# Identifying Brain Regions Related to Word Prediction During Listening to Japanese Speech by Combining a LSTM Language Model and MEG

**DOI:** 10.1101/2021.03.25.436887

**Authors:** Yuta Takahashi, Yohei Oseki, Hiromu Sakai, Michiru Makuuchi, Rieko Osu

## Abstract

Recently, a neuroscientific approach has revealed that humans understand language while subconsciously predicting the next word from the preceding context. Most studies on human word prediction have investigated the correlations between brain activity while reading or listening to sentences on functional magnetic resonance imaging (fMRI) and the predictive difficulty of each word in a sentence calculated by the N-gram language model. However, because of its low temporal resolution, fMRI is not optimal for identifying the changes in brain activity that accompany language comprehension. In addition, the N-gram language model is a simple computational structure that does not account for the structure of the human brain. Furthermore, it is necessary for humans to retain information prior to the N-1 word in order to form a contextual understanding of a presented story. Therefore, in the present study, we measured brain activity using magnetoencephalography (MEG), which has a higher temporal resolution than fMRI, and calculated the prediction difficulty of words using a long short-term memory language model (LSTMLM), which is based on a neural network inspired by the structure of the human brain and has longer information retention than the N-gram language model. We then identified the brain regions involved in language prediction during Japanese-language speech listening using encoding and decoding analyses. In addition to surprisal-related regions revealed in previous studies, such as the superior temporal gyrus, fusiform gyrus, and temporal pole, we also found relationships between surprisal and brain activity in other regions, including the insula, superior temporal sulcus, and middle temporal gyrus, which are believed to be involved in longer-term, sentence-level cognitive processing.

## 1. Introduction

In recent years, a neuroscientific approach has revealed that humans are always predicting what will happen next (Clark, 2013). This principle also applies to language comprehension, a process during which humans understand language while subconsciously predicting the next word from the preceding context (DeLong et al., 2005; Dikker et al., 2010).

In recent neuroscientific research of human word prediction, experiments have been conducted under natural conditions of reading (Wehbe et al., 2014) or listening to (Brennan et al., 2012; Willems et al., 2016) stories or speeches instead of under strictly controlled conditions with intermittent visual stimuli. Most studies have investigated the correlation between brain activity while reading or listening to sentences on functional magnetic resonance imaging (fMRI) and the predictive difficulty of each word in a sentence determined by the N-gram language model (Russo et al., 2020; Willems et al., 2016). However, because of its low temporal resolution, fMRI is not optimal for identifying the changes in brain activity accompanying language comprehension. In particular, it is more difficult to assess brain activity during comprehension of languages that consist of short words, such as Japanese, using fMRI than languages with longer words, such as English or other European languages. Methods that measure neuroelectrical activity, including magnetoencephalography (MEG) and electroencephalography (EEG), have higher temporal resolutions than fMRI, and are more suitable for detecting activity related to language processing in an ecologically valid context.

The N-gram language model has been used for natural language processing, including machine translation, and is known for its high accuracy and low computational cost. This model is also often used in neurolinguistic research to study language processing in the human brain. However, the N-gram language model involves a simple computational structure that does not account for the structure of the human brain, implying it may be inappropriate to use in this context. Furthermore, the N-gram language model calculates the predictive difficulty of each word based on information from the word, N-1, which occurs before the word one before. Humans, however, must retain information from prior to the N-1 word in order to form a contextual understanding of a presented story. Therefore, it is questionable if this model has validity for evaluating language processing in the human brain.

In contrast, the long short-term memory language model (LSTMLM) is based on a neural network inspired by the structure of the human brain, and it can be used to calculate the difficulty of prediction based on long-term past information. Therefore, the LSTMLM may be more suitable than the N-gram language model for understanding the human brain during neurolinguistic research.

In the present study, we measured brain activity using MEG, which has a higher temporal resolution than fMRI, and calculated the prediction difficulty using the LSTMLM, which retains information longer than the N-gram language model. We then identified the brain regions involved in language prediction during Japanese-language speech listening using encoding and decoding analyses (Fig. 1). To our knowledge, this is the first study to use MEG and the LSTMLM to investigate language processing in the human brain.

**Fig. 1.**
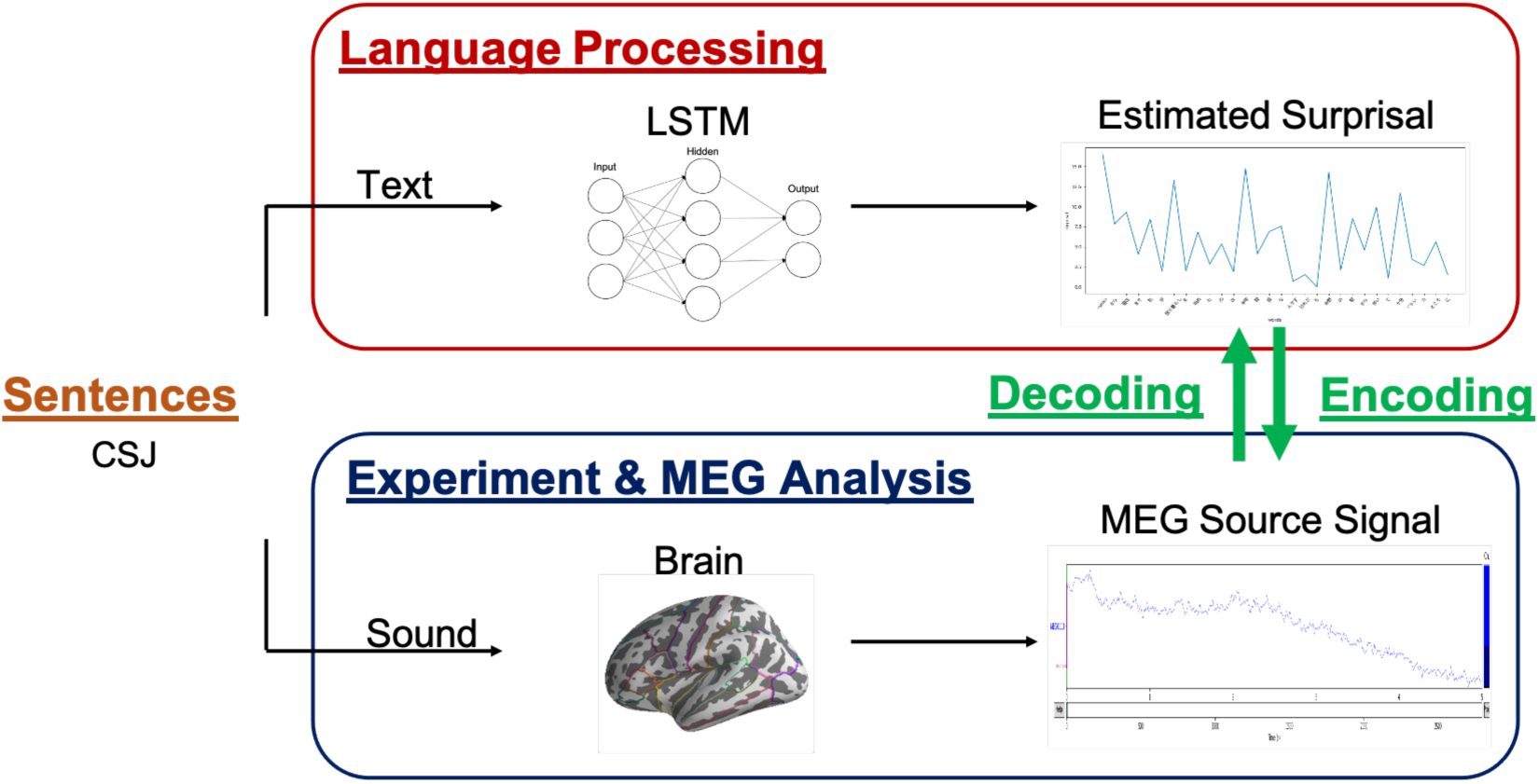
Experimental and data analysis design. We used 201 Japanese speeches from a database called the Corpus of Spontaneous Japanese (CSJ). For the magnetoencephalography (MEG) experiment, the 10 study participants listened to four of these speeches. Using this data, the source signal while listening to speeches was estimated. For language processing, the transcripts of each speech were first divided into long unit words (LUWs). The LUWs of the remaining 197 speeches were then inputted into the long short-term memory language model (LSTMLM) as training data, while the four speeches used in the MEG experiments were inputted as test data, with estimation of the surprisal for each LUW of the speeches used in the experiments. The relationships between MEG source signals and surprisal values were then investigated using encoding and decoding analyses. LSTM, long short-term memory

### 1.1 Language Model

The N-gram language model, which has been widely used in previous studies, is based on the Markov assumption that the probability of a word’s occurrence does not depend on the preceding word but only on the most recent N-word. The probability of the *t*-th word occurrence *p*(*w_t_*) is expressed by the following formula:

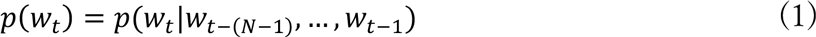

Although the N-gram language model has a low computational cost and high accuracy, it is difficult to account for the meaning of sentences and the relationships between words using this model because each computation is independent. In addition, unknown word combinations not included in the training data cannot be calculated by the N-gram language model; therefore, a large amount of data needs to be trained.

Recently, an LSTMLM (Sundermeyer et al., 2012), which uses long short-term memory (LSTM) with a recurrent neural network (RNN) architecture, has been introduced as a practical language model for artificial intelligence research. LSTMLM has a recursive structure and predicts the next word by representing words as vectors and continuously synthesizing the information from one step earlier in the hidden layer of each step. Therefore, in principle, the information from all preceding words is kept chronologically, and the probability of the occurrence of a word is calculated based on this long-term information. LSTMLM is generally said to be superior to the N-gram language model for word prediction accuracy and adaptation to unknown words (Jozefowicz et al., 2016). Furthermore, when humans understand a sentence while subconsciously predicting the next word, they must incorporate information not only from the preceding few words but also from entire sentences that appeared previously; that is, in context. Therefore, the LSTMLM may be a more human-like model than the N-gram language model.

### 1.2. Surprisal

One indicator of a word’s prediction difficulty is its surprisal. The surprisal is a measure of the unexpectedness of a target word, and it is used in machine translation and other natural language processing applications. The surprisal of the *t*-th word is calculated using the following formula:

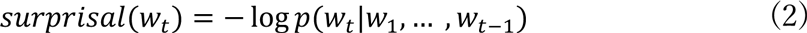

This formula indicates that when the probability of the occurrence of an observed word is high (easy to predict), the surprisal is low, and, when the probability of the occurrence of an observed word is low (difficult to predict), the surprisal is high. The surprisal is considered to be a measure of the context-dependent cognitive load (Hale, 2001; Levy, 2008), and it has been found to correlate with the amplitude of the event-related potential (ERP) of MEG during sentence comprehension (Lau et al., 2009).

### 1.3. Encoding and Decoding Analyses

In this study, we conducted two types of analyses: encoding and decoding. These analyses are both designed to reveal the role of specific brain regions in language processing; however, they differ in the information that they target (Naselaris et al., 2011). Encoding analysis predicts brain activity from external stimuli or behavior, revealing how this activity is represented in each voxel. On the other hand, decoding analysis predicts external stimuli or behavior from brain activity, thereby revealing the roles of specific brain regions by examining the information represented by the activity of multiple voxel patterns.

It is important to analyze brain mechanisms using approaches that target different types of information. For example, in the field of vision research, one study used encoding analysis to predict fMRI activity while looking at a specific image (Kay et al., 2008), while another study used decoding analysis to identify the direction of a stimulus at which the subject was looking based on fMRI activity patterns in the primary visual cortex (Kamitani and Tong, 2005). In word comprehension research, one study used encoding analysis to predict fMRI activity while reading a specific word (Mitchell et al., 2008), while another study used decoding analysis to reveal the congruency of fMRI activity patterns while bilingual listeners listened to the same word across different languages (Correia et al., 2014). While encoding analysis has primarily been used in previous MEG studies of language processing (Gwilliams et al., 2016), decoding analysis has been used less frequently.

Therefore, in this study, we conducted both an encoding analysis to predict MEG signals from the surprisal and a decoding analysis to predict the surprisal from MEG signals.

## 2. Methods

### 2.1. Participants and Data Acquisition

We recruited 13 healthy participants for this study; however, one of these participants was excluded because he fell asleep during the experiments, and another two were excluded because of improper measurements. Therefore, we included data from 10 participants (five females, mean age 20.89±1.9 years [M ± SD]) for further analysis. Written informed consent was obtained from all participants, and the study was approved by the Ethics Committee of the National Rehabilitation Center for Persons with Disabilities in Japan.

We collected MEG data with a 204-channel axial gradiometer and a 102-channel axial magnetometer system (Elekta Ltd., Helsinki, Finland) at the National Rehabilitation Center for Persons with Disabilities in Japan. The signals were measured at a sampling frequency of 1000 Hz. At the time of measurements, three markers (nasion and left and right canals) were attached to the participant’s head to determine the position of the head with respect to the MEG sensor.

### 2.2 Experimental Procedure and Stimulus Materials

For these experiments, participants were asked to listen to four speeches while brain activity was measured using MEG. These four speeches were selected from 201 manually labeled Japanese speeches from the “Corpus of Spontaneous Japanese (CSJ)” (Maekawa, 2003) created by the National Institute for Japanese Language and Linguistics. These speeches did not overlap in theme and consisted of a fast female speech, a slow female speech, a fast male speech, and a slow male speech. Each speech was approximately 10 min in length (12:46, 8:41, 11:14, and 10:18). While in the MEG scanner, participants listened to the speeches via earphones and were instructed to simultaneously look at a blank screen. The four speeches were presented in a random order to each participant, and the participants were allowed to take breaks inside the MEG scanner between speeches. Participants were informed in advance that they would be asked to answer five simple comprehension questions after listening to each speech and were required to listen carefully to the speeches to understand their content. The comprehension questions were presented as text on the screen, and participants were required to indicate whether the questions were correct or incorrect based on the speech content by pushing buttons on the response pad with the middle and index fingers of their left hand.

### 2.3. Language Processing

We estimated the number of words that appeared in the four speeches used in the MEG experiments. The transcript of each speech was divided into long unit words (LUWs), which involves the division of phrases into content and function words, with the surprisal predicted for each LUW. We estimated surprisal using the neural language model proposed by Van Schijndel and Linzen (2018), which is based on LSTM. This model consists of an embedding layer, two hidden layers, and an output layer, with 200 units for each layer (Fig. 2). The LUWs were vectorized in the embedding layer and inputted into the hidden LSTM layer. In this hidden LSTM layer, information from the hidden layer one step earlier and input information from that point were combined and sent to the output layer of that step, as well as to the hidden layer of the next step to calculate the hidden state. In the output layer, the softmax function computed the probability of the occurrence of the next LUW. We used cross-entropy for loss function and perplexity, the inverse of probability, to evaluate the model. The following parameters were set to default values: the number of steps for backpropagation through time (BPTT) was set to 35, the batch size was set to 20, the learning rate was set to 20, and the dropout ratio was set to 0.2. The epoch size for each piece of training data was set to 40; however, training was stopped early if the validation loss remained the same for three consecutive epochs.

**Fig. 2.**
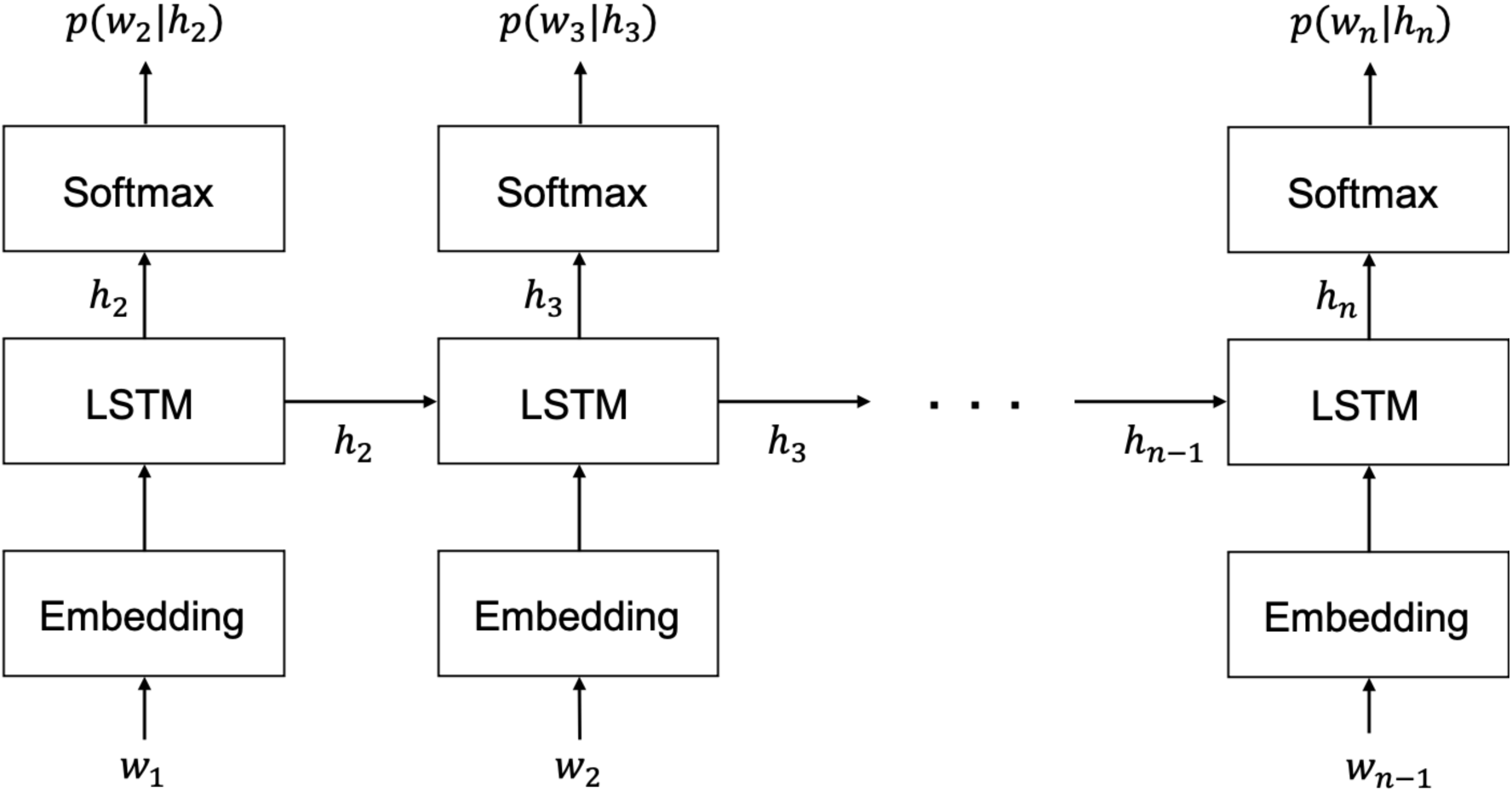
Structure of the long short-term memory language model (LSTMLM). *W_n_*_-1_ is the (*n*-1)-th long unit word (LUW), and each LUW is converted to a vector in the embedding layer and then inputted to the hidden layer, which is then combined with the hidden state *h_n_*_-1_ sent from the (*n*-2)-th hidden layer, and the hidden state *h_n_* is calculated. The hidden state *h_n_* is sent to both the (*n*-1)-th output layer and the *n*-th hidden layer. In the output layer, the probability *p*(*w_n_*|*h_n_*) of the *n*-th LUW occurring up to the (*n*-1)-th LUW is calculated using the softmax function.

For training, the 197 CSJ speeches not used in the MEG experiments were utilized. These speeches were transcribed and divided into LUWs in the same manner as the speeches used in the MEG experiments. Of these, 95% (187 speeches; 351,018 LUWs in total) were used as training data, and the remaining 5% (10 speeches; 18,330 LUWs in total) were used as validation data. Each of the four speeches used in the MEG experiments were then entered into the trained model, and the surprisal per LUW of each speech was estimated.

For comparison, we also attempted to train a trigram language model, or a language model with three-word N-grams, using the same data.

### 2.4. MEG Data Preprocessing

We used Python 3.7 and the Eelbrain module (https://pythonhosted.org/eelbrain/) for preprocessing and analysis of the MEG data according to the Eelbrain pipeline. For all raw data, a Maxwell filter (Taulu and Kajola, 2005) was applied to remove magnetic noise, and a 1‒40-Hz bandpass filter was applied. We then performed an independent component analysis (ICA) of data from each speech and each participant and reconstructed the signal, leaving only those components with a cumulative explained variance of less than 99%.

In order to create epochs, we identified the onset of each LUW from the speech data. Each epoch was determined to start at 200 ms after the onset of each LUW, with a window size of 400 ms. As a result, the total number of epochs in the four speeches was 7,966. The data in each epoch were downsampled to 100 Hz; therefore, the number of bins in each epoch was 40.

Next, we estimated the source signal. To reconstruct the position of the MEG sensor relative to the FreeSurfer average brain (CorTechs Labs Inc., Lajolla, CA), the transformation from the marker positions of each participant to the reference points of the average brain were applied to the MEG sensor position. We then created an ico-4 source space consisting of 5,124 vertices and calculated the activity at each vertex, i.e., the source signal, using noise normalization with the dynamical Statistical Parametric Mapping (dSPM) method (Dale et al., 2000). Orientation was constrained to only estimate the current flow orthogonal to the cortical surface. Therefore, the sign of the estimates indicated the direction of the current relative to the cortical surface, and a positive sign represented the signal from the source to the cortical surface.

### 2.5. Encoding Analysis

We regressed the MEG source signals from the surprisal estimated using the LSTMLM and estimated the brain regions correlated with the magnitude of the surprisal. We ran spatiotemporal permutation cluster tests (Maris and Oostenveld, 2007) using the TwoStageTest in Eelbrain. The spatiotemporal permutation cluster tests consisted of three stages (Fig. 3). In the first stage, the MEG source signal sequences of the 7,966 epochs were fitted by a linear regression model for each subject at each of the 5,124 vertices and at each of the 40 time points within the epoch, with 7,966 corresponding surprisal value sequences as independent variables using ordinary least squares. As a result, the regression coefficient, beta, at each source and at each time point was obtained for each participant. In the second stage, 10 beta values at each source and at each time point for all participants were tested using the one-sample *t*-test to determine if they were significantly different from zero. This analysis resulted in a *t*-value and *p*-value for each source and each time point, which were then used to construct a 5124 × 40 spatiotemporal map for each subject. In the third stage, a permutation cluster test was run. First, a cluster was defined as a group of points on the map whose spatiotemporally adjacent points had *t*-values with a *p*-value of < .05. The points in a cluster were required to have *t*-values with the same polarity, and clusters with different polarities were identified as separate regions. If the cluster consisted of at least 10 sources with two consecutive time points, the *t*-values within that cluster were summed to yield the cluster statistics. For comparison with these cluster statistics, permutation statistics were calculated for each cluster by randomly selecting the same number of points as that cluster from the 5124 × 40 spatiotemporal map and summing the *t*-values of these points. This process was repeated 10,000 times for that cluster, and the number of times the cluster statistic was below the permutation statistic, divided by 10,000, was defined as the corrected *p*-value for that cluster. Clusters with a corrected *p*-value of < .05 were considered to be significant clusters.

**Fig. 3.**
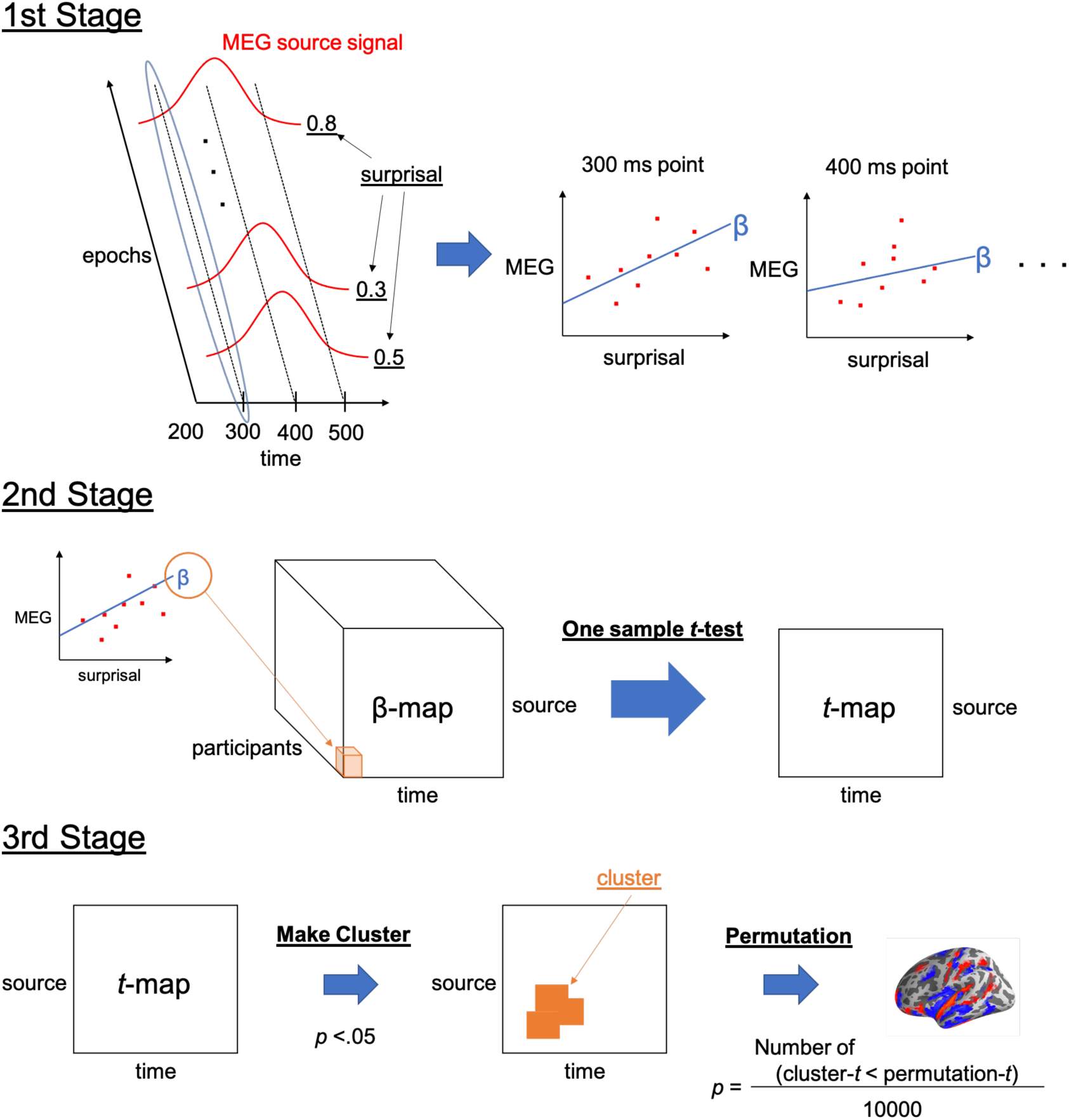
Summary of spatiotemporal permutation cluster tests. These tests consisted of three stages. In the first stage, the regression coefficient beta between the surprisal and the magnetoencephalography (MEG) source signal at each source and at each time point were obtained for each participant. In the second stage, a one-sample *t*-test was performed to determine whether a subject’s beta was different from zero at each time point and at each source, and a *t*-map was created. In the third stage, spatiotemporally adjacent points with *p*-values of < .05 on this *t*-map were combined to form clusters. Then, for each cluster, cluster-t, which was the sum of the *t*-values in the cluster, and the permutation-t, which was the sum of the *t*-values randomly extracted from the same number of points in the cluster, were compared 10,000 times as a permutation *t*-test. The number of times that the permutation-t was greater than the cluster-t divided by 10,000 was defined as the corrected *p*-value for that cluster.

### 2.6. Decoding Analysis

We estimated the brain regions responsible for surprisal-related information processing using the decoding method. That is, we reconstructed the data from multiple MEG source signals and identified regions with high regression accuracy. For the decoding analysis, the MEG source signals were downsampled to 50 Hz after preprocessing. Therefore, the data in each 400-ms epoch, which started 200 ms after LUW onset, had 20 time points. The cortex was then divided into 68 regions based on the ‘Desikan-Killiany’ cortical atlas (Desikan et al., 2006), and each region was decoded separately. For regression, we used the LASSO regression model (alpha = 0.01), which is a linear model with the addition of an L1 regularization term. Scikit-learn (https://scikit-learn.org), a Python library for machine learning, was used for standardization and model fitting. The root mean square error (RMSE) was used to evaluate the model. Regression of the surprisal at each time point in the epoch was performed for each region, and the features were defined as the activity of all MEG sources at the time point included within the corresponding region. For each region, the activity was standardized over all sources, epochs, and time points to have a mean of 0 and a variance of 1. Cross-validation was repeated five times, with 80% of the 7,966 epochs used as training data and 20% used as test data, with the average of five RMSE values considered to be the regression accuracy. The integral value calculated by the LSTM was set as the training label for the regression model. The regression accuracies were then averaged for every 100 ms (every five time points). For comparison, surprisal labels were randomized among epochs, and the surrogate regression accuracies were calculated. These surrogate accuracies were then averaged every 100 ms. This process was repeated 100 times for each cross-validation for a total of 500 iterations.

To identify regions where the regression accuracy of the original data was significantly higher than the regression accuracy of the surrogate data, we performed two statistical analyses. The first was a paired-samples *t*-test that compared the mean accuracy of 500 pieces of surrogate data with the regression accuracy of the original data for the 10 participants in each region to determine if the original regression accuracy was significantly higher than the mean surrogate accuracy. Multiple comparisons were corrected by the false discovery rate (FDR) using the Benjamini-Hochberg (BH) method (Benjamini and Hochberg, 1995), and regions with a corrected *p*-value of < .01 were defined as having a significant difference. The second statistical analysis was a one-sample *t*-test to determine whether the actual accuracy was significantly different than the surrogate accuracy distribution for each region in each participant. The BH method was used to correct for multiple comparisons. A region where greater than 90% of participants had a corrected *p*-value of < .05 was defined as having a significant difference. Regions that demonstrated significance using both of these statistical analyses were defined as having significantly higher regression accuracy.

## 3. Results

### 3.1. Language Processing

The LSTM training was completed in the 9th epoch. The value of the validation perplexity was 137.00. We independently inputted each of the four speeches used in the experiment into this trained model and predicted the surprisal for each LUW (Fig. 4). The average magnitude and standard deviation (SD) of the surprisal was 6.97±5.51.

**Fig. 4.**
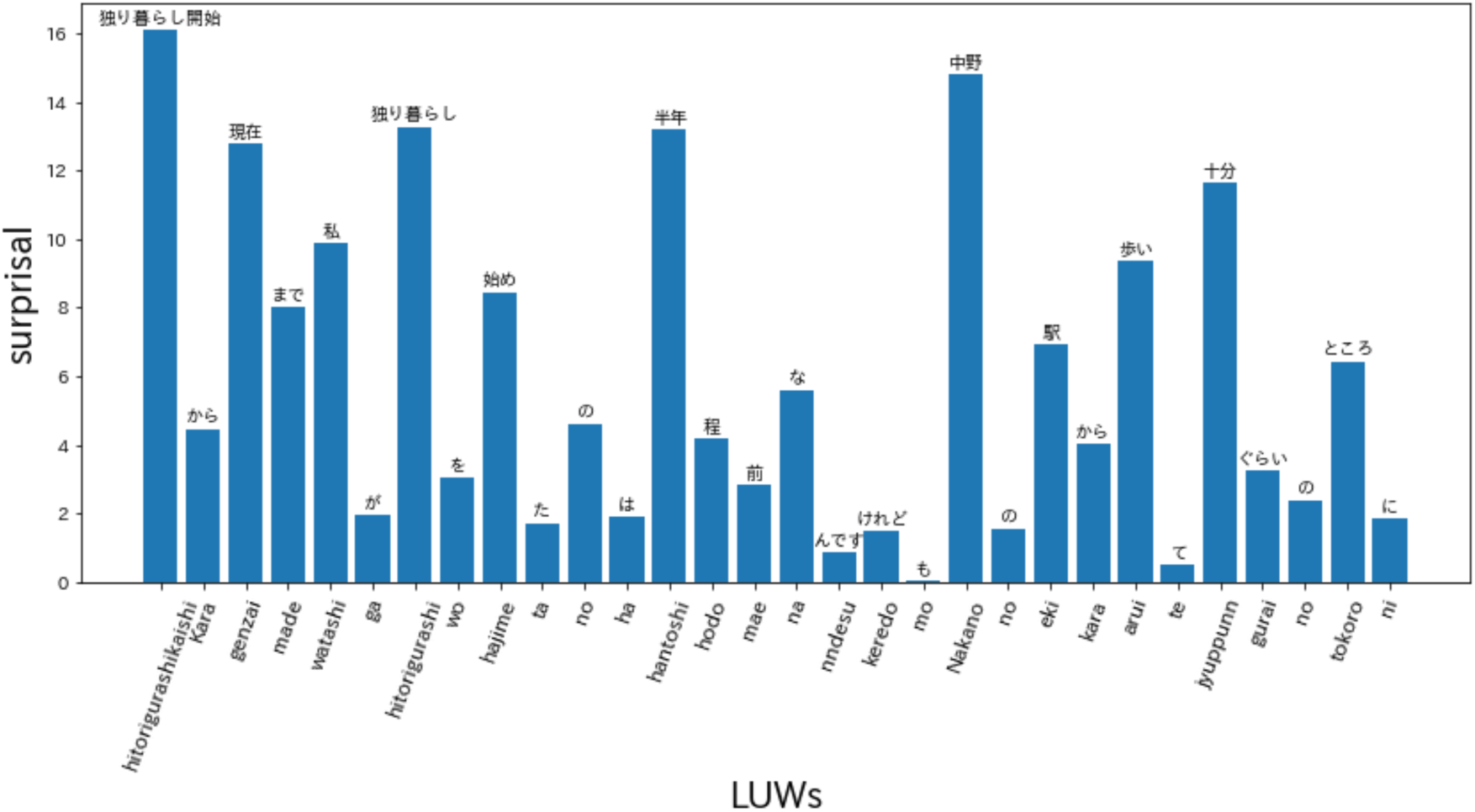
Example surprisal prediction results using the long short-term memory language model (LSTMLM). We predicted surprisal for each long unit word (LUW) for the four speeches used in the experiments. The average magnitude and standard deviation of the surprisal was 6.97±5.51.

We also attempted to train the trigram language model using the same data from the 197 speeches; however, the validation perplexity value was 230666.94; that is, learning did not converge because of the small data size.

### 3.2. Encoding Analysis

In the encoding analysis, 145 clusters were formed (Fig. 5). Among these, three clusters showed corrected *p*-values less than 1% in the permutation *t*-test, which examined the correlation between the surprisal values and MEG source activity. The first of these clusters contained a part of the left insula; the left supramarginal gyrus (SMG); the left temporal pole; the region around the left superior temporal gyrus (STG), including the transverse temporal gyrus (TTG); and the region around the left fusiform gyrus (FuG), including the entorhinal cortex and the parahippocampal gyrus (PHG) (corrected *p* < .005) (Fig. 6-A). The second was a left hemispheric cluster containing the region around the left inferior frontal gyrus (IFG), including the pars opercularis, pars triangularis, pars orbitalis, and the orbitofrontal cortex (OFC), as well as the left frontal operculum (FOP), which is adjacent to the insula and located ventrally and medially to the pars opercularis and pars triangularis (corrected *p* < .005) (Fig. 6-B). The third was a right hemispheric cluster contralateral to the second cluster containing the right IFG and the right FOP (corrected *p* < .005) (Fig. 6-C). In these strongly correlated regions, the peak correlation occurred at 300 ms after the start of the LUW in each case.

**Fig. 5.**
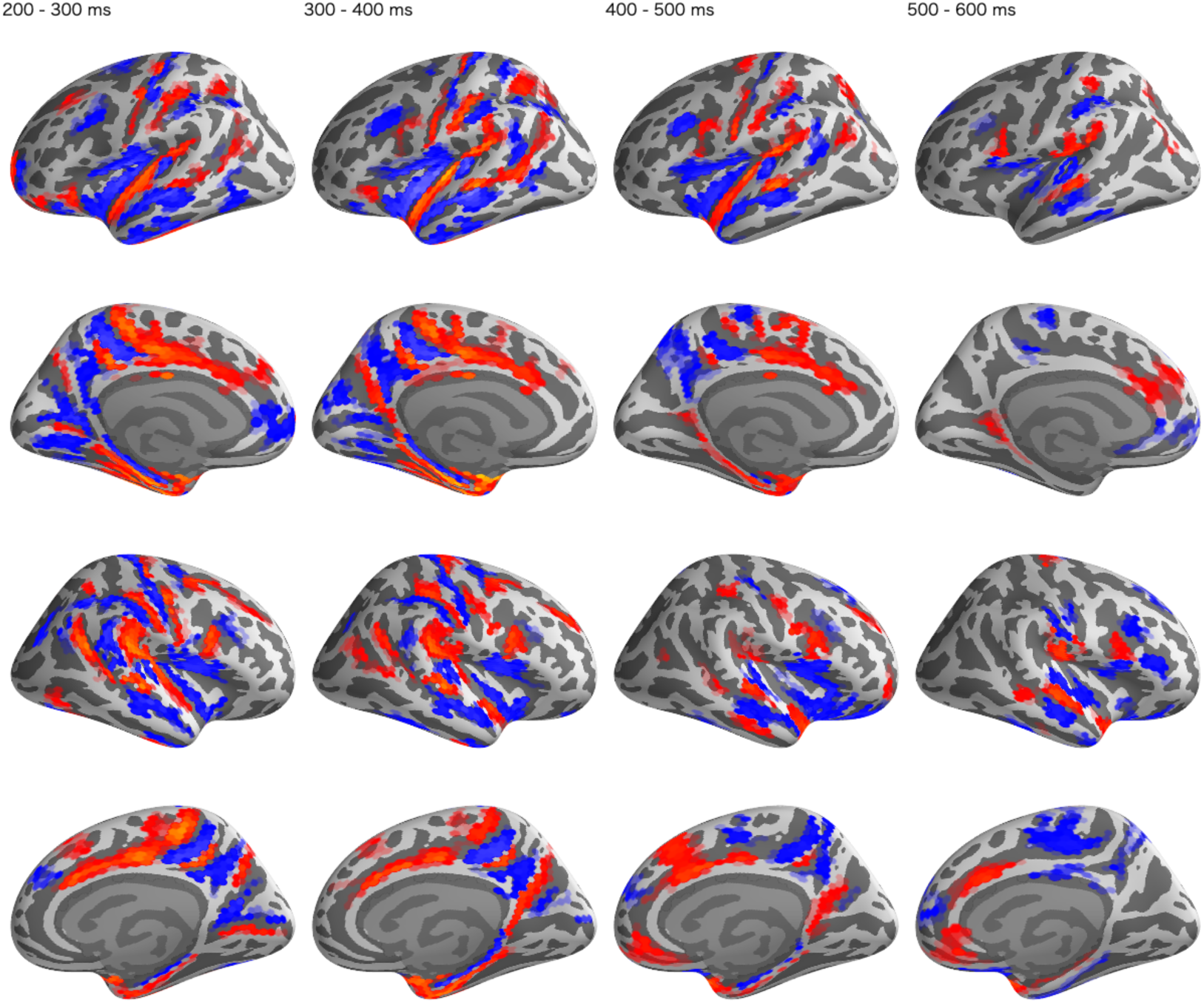
Spatiotemporal maps of clusters formed in the encoding analysis. In total, 145 clusters were generated. The red regions are clusters with positive correlations between surprisal values and magnetoencephalography (MEG) source activities, and the blue regions are those with negative correlations.

**Fig. 6.**
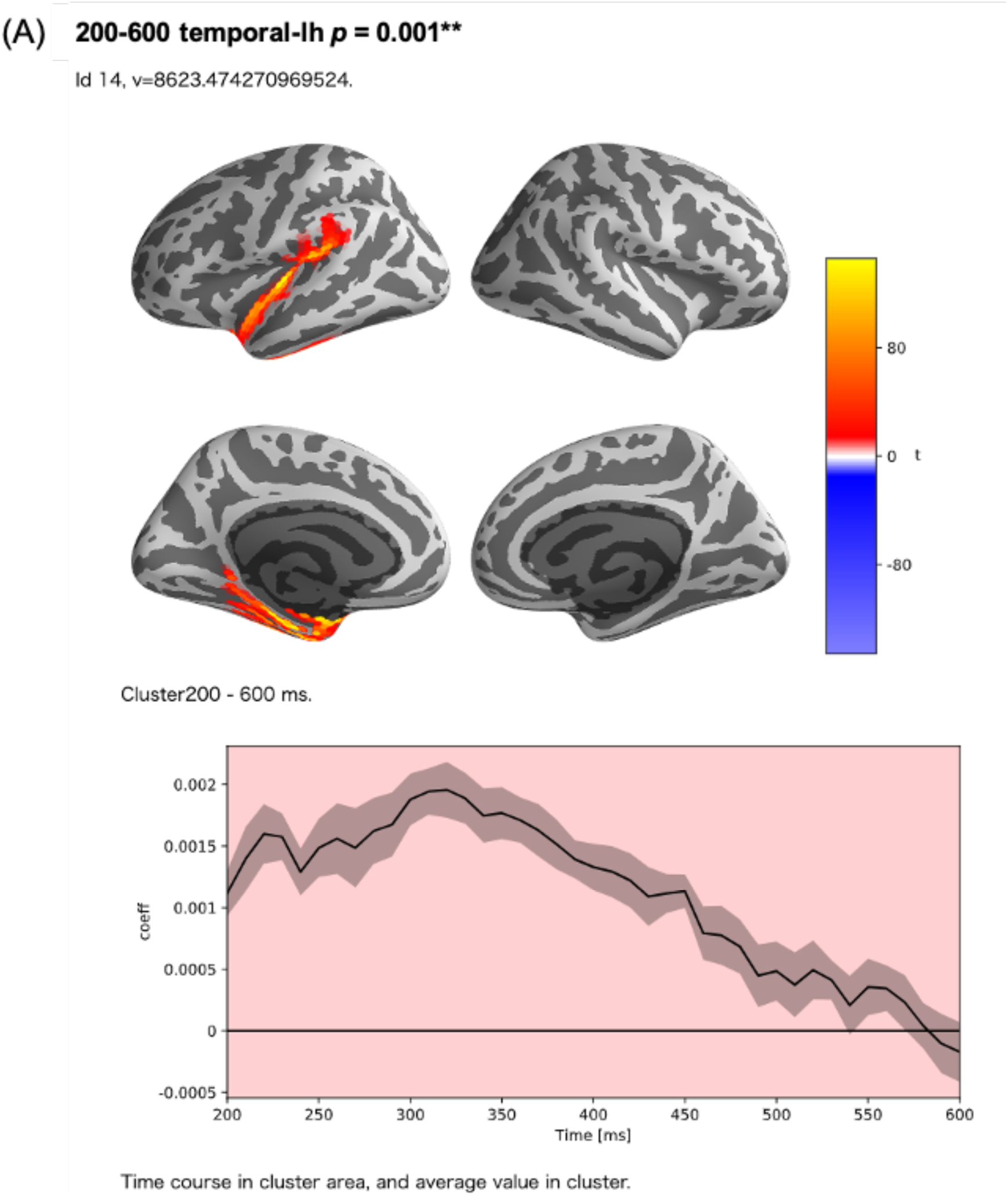
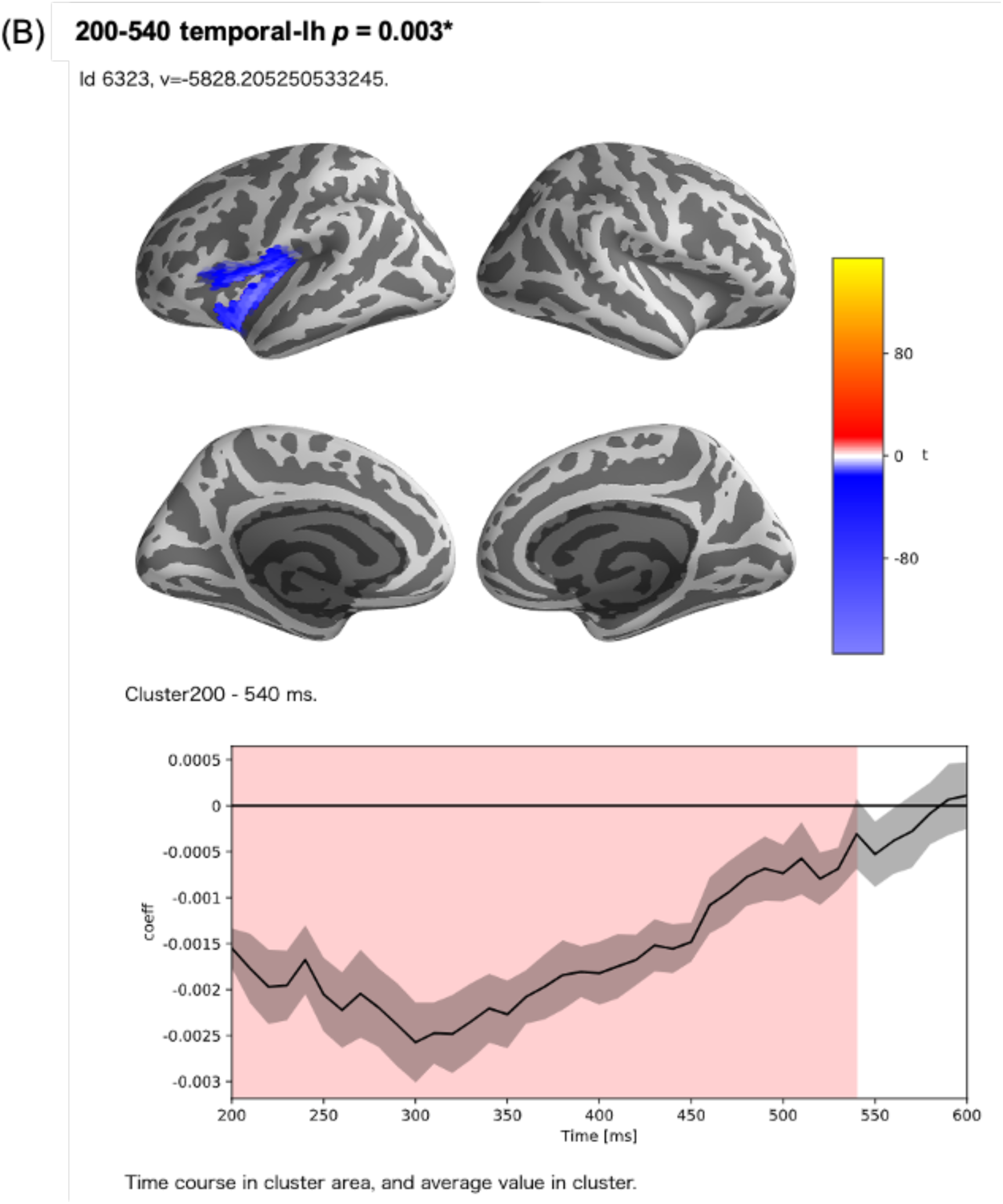
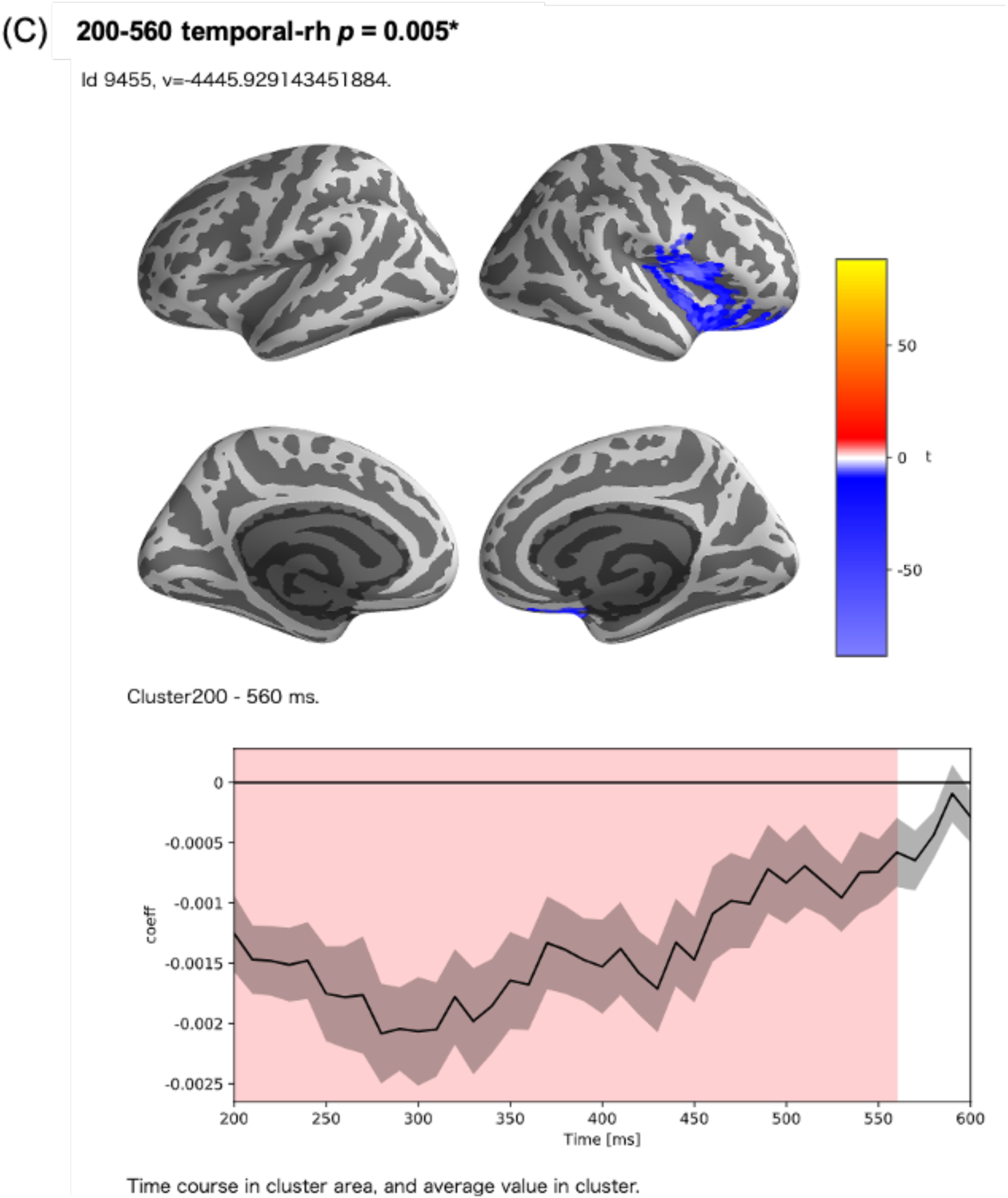
Clusters with a corrected *t*-value less than 1% in the permutation *t*-test. (A) A left hemispheric cluster generated with 216 sources and a time range of 200‒600 ms after the onset of LUWs. In this cluster, there was a strong positive correlation (cluster *t*-value = 8623.47, corrected *p*-value = 0.001). This cluster consisted of a part of the left insula, the left supramarginal gyrus (SMG), the left temporal pole, the region around the left superior temporal gyrus (STG), and the region around the left fusiform gyrus (FuG). (B) A left hemispheric cluster generated with 128 sources and a time range of 200‒540 ms after the onset of LUWs. In this cluster, there was a negative correlation (cluster *t*-value = −5828.21, corrected *p*-value = 0.003). This cluster consisted of the region around the left inferior frontal gyrus (IFG) and the left frontal operculum (FOP). (C) A right hemispheric cluster generated with 169 sources and a time range of 200‒560 ms after the onset of LUWs. In this cluster, there was a negative correlation (cluster *t*-value = −4445.93, corrected *p*-value = 0.005). This cluster consisted of the region around the right IFG and the right FOP.

### 3.3. Decoding Analysis

In the encoding analysis, there were many regions where correlations peaked at 300 ms after the start of a LUW and disappeared between 500 and 600 ms. Therefore, for the decoding analysis, we analyzed three time ranges, 200‒300 ms, 300‒400 ms, and 400‒500 ms after the onset of a LUW.

First, decoding analysis was performed for the 200‒300-ms time range. A paired-samples *t*-test was conducted for all participants to compare whether the regression error of the original data was significantly smaller than the mean regression error of the surrogate data, with the left insula, left medial orbital frontal cortex (mOFC), left SMG, left STG, and right isthmus-cingulate cortex (ICC) showing significance at the 1% level (Fig. 7-A, B). Additionally, a one-sample *t*-test was conducted to compare whether the regression error of the original data was significantly smaller than the regression error of the surrogate data for each subject. The bilateral insula, bilateral postcentral gyrus (PoG), left Banks superior temporal sulcus (BanksSTS), left STG, left SMG, and right precentral gyrus (PrG) (Fig. 7-C, D) were found to have significant differences in more than nine participants (90% of total participants). Regions with significant differences using both of these statistical analyses were defined as having a significantly higher regression accuracy in that time range. Therefore, for the time range of 200‒300 ms, three regions, including the left insula, left STG, and left SMG, were determined to have significantly higher regression accuracies (Fig. 10-A).

**Fig. 7.**
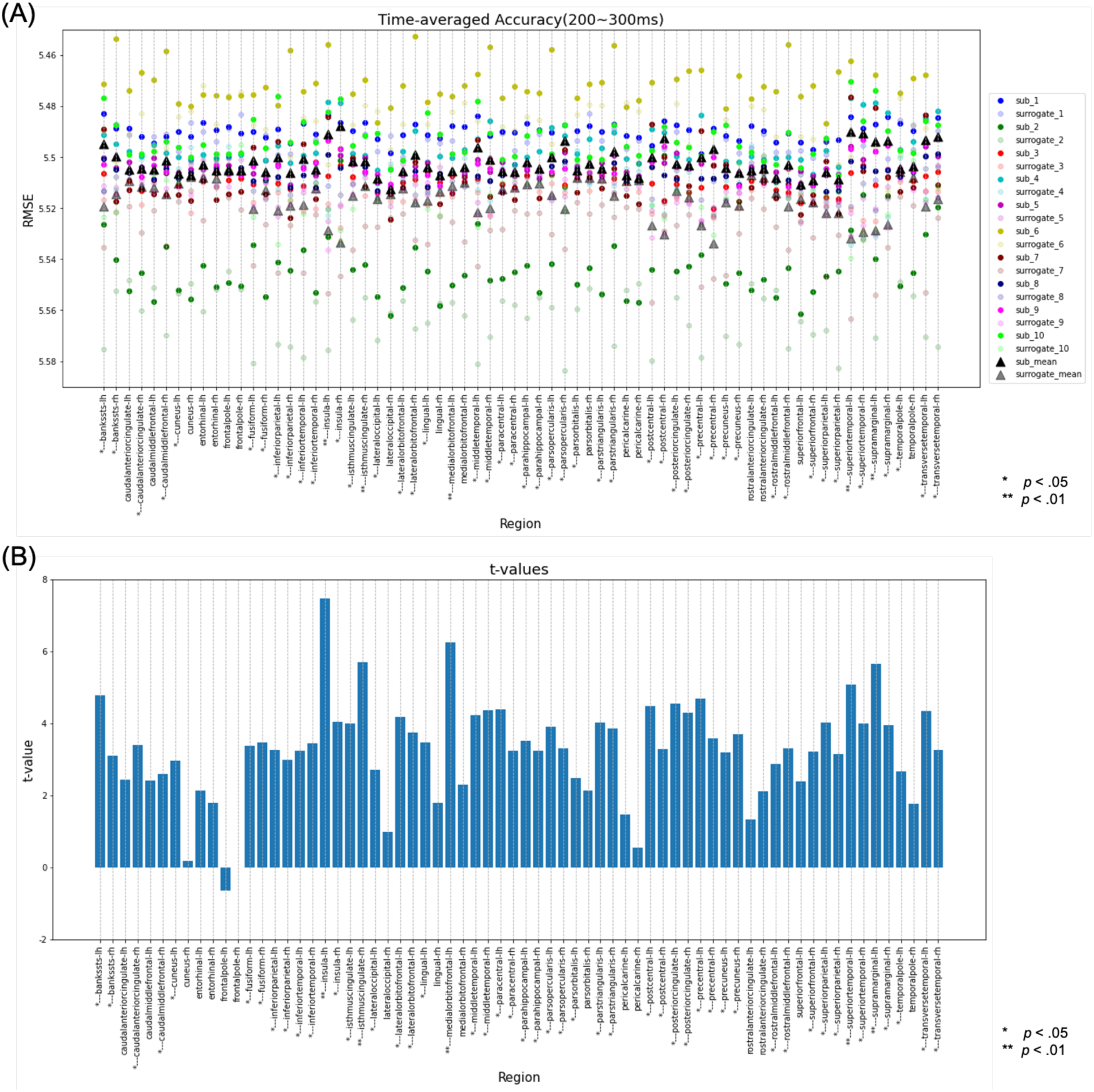
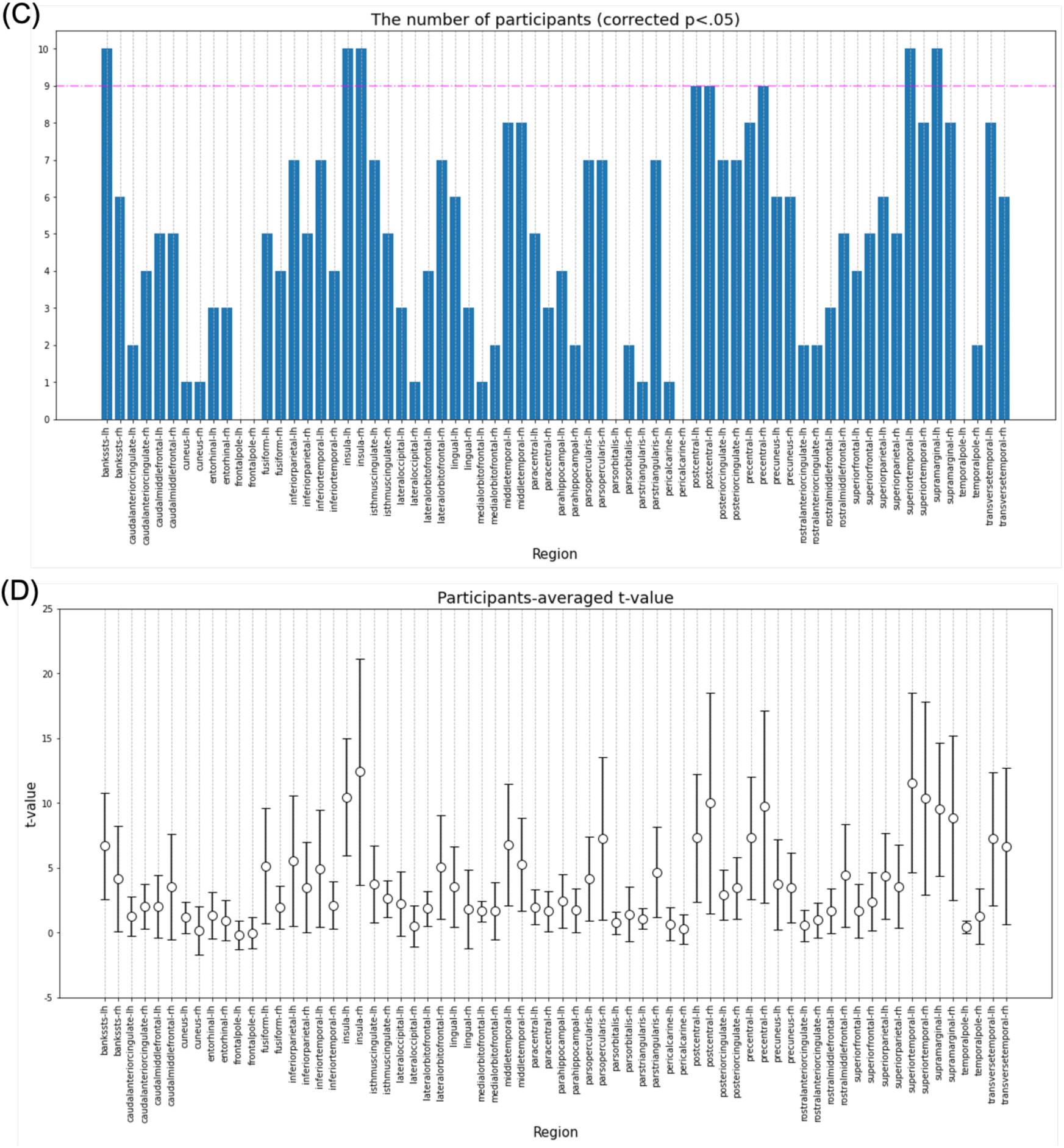
Results of the decoding analysis for each region in the 200‒300-ms time range. (A) The regression error of the original data and the mean regression error of the surrogate data in the time range of 200‒300 ms was plotted for each subject. The paired-samples *t*-test for all subjects compared whether the regression error of the original data was significantly smaller than the mean regression error of the surrogate data. *regions with a corrected *p*-value of < .05 and ** regions with a corrected *p*-value of < .01. (B) The *t*-value for each region in the paired-samples *t*-test. (C) Number of subjects with a corrected *p*-value of < .05 on the one-sample *t*-test comparing whether the regression error of the original data was significantly smaller than the regression error of the surrogate data for each subject. The regions where more than 90% of subjects (above the pink line) had a corrected *p*-value of < .05 were defined as regions with significant differences in the one-sample *t*-test. (D) Subject mean and variance of *t*-values in the one-sample *t*-test.

Next, decoding analysis was performed for the 300‒400-ms time range. In this analysis, the bilateral insula, bilateral PHG, bilateral PoG, bilateral posterior cingulate cortex (PCgG), bilateral PrG, left BanksSTS, left lingual gyrus (LgG), left middle temporal gyrus (MTG), left paracentral lobule (PCL), left STG, left SMG, left TTG, right caudal anterior cingulate cortex (CaudalACC), right FuG, right pars triangularis, and right rostral middle frontal gyrus (RostralMFG) (Fig. 8-A, B) were found to be significantly different using the paired-samples *t*-test. Additionally, the bilateral insula, bilateral PrG, bilateral STG, left BanksSTS, left meddle temporal, left SMG, and left TTG were found to have significant differences in more than nine participants using the one-sample *t*-test (Fig. 8-C, D). Therefore, the regions with significant differences using both of these statistical analyses were the bilateral insula, bilateral PrG, bilateral STG, the left BanksSTS, the left MTG, the left SMG, and the left TTG (Fig. 10-B).

**Fig. 8.**
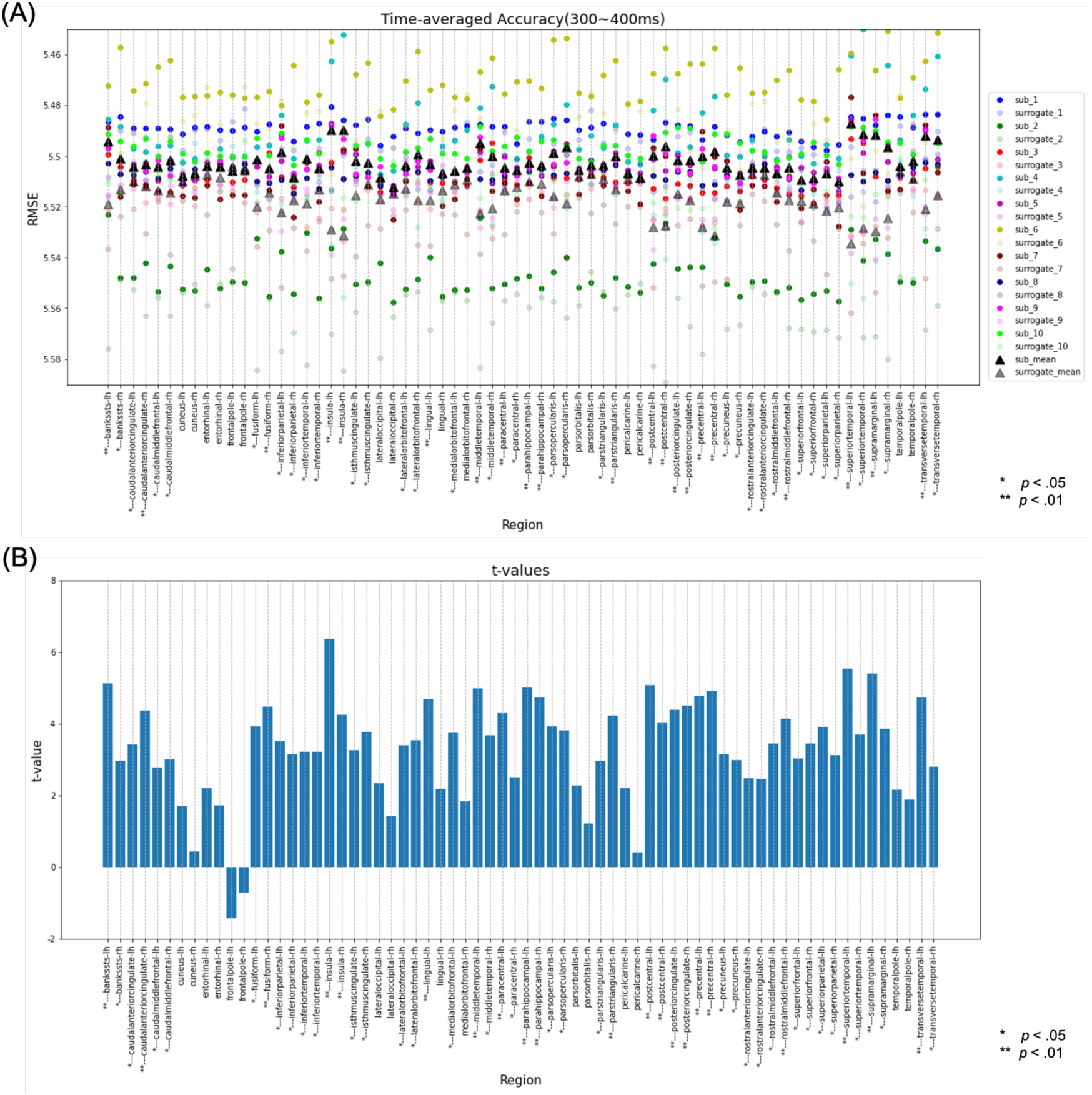
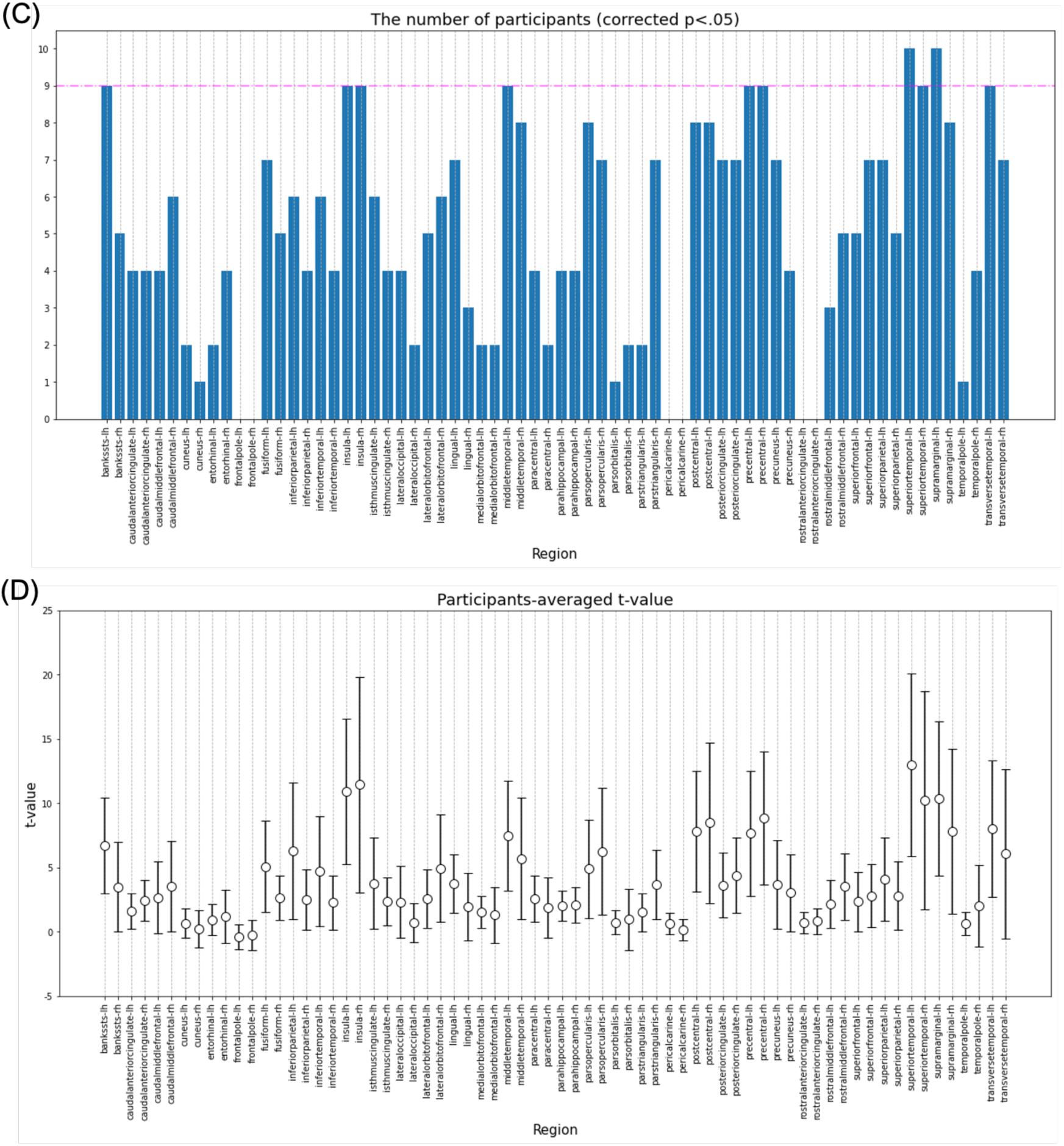
Results of the decoding analysis for each region in the time range of 300–400 ms. (A) The regression error of the original data and the mean regression error of the surrogate data in the time range of 300‒400 ms were plotted for each subject. The paired-samples *t*-test for all subjects compared whether the regression error of the original data was significantly smaller than the mean regression error of the surrogate data. *Regions with a corrected *p*-value of < .05 and ** regions with a corrected *p*-value of < .01. (B) The *t*-value for each region in the paired-samples *t*-test. (C) The number of subjects with a corrected *p*-value of < .05 in the one-sample *t*-test comparing whether the regression error of the original data was significantly smaller than the regression error of the surrogate data for each subject. Regions where more than 90% of the total subjects (above the pink line) had a corrected *p*-value of < .05 were defined as regions with significant differences in the one-sample *t*-test. (D) Subject mean and variance of *t*-values in the one-sample *t*-test. RMSE, root mean square error.

Next, decoding analysis was performed for the 400‒500-ms time range. In this analysis, the left BanksSTS, left PoG, left superior parietal lobule (SPL), left STG, left SMG, right PrG, and right RostralMFG were found to have significant differences using the paired-samples *t*-test (Fig. 9-A, B). However, in the one-sample *t*-test, no regions were found to have significant differences in more than nine participants (Fig. 9-C, D). Therefore, there was no region where both of these statistical analyses showed significant differences, indicating there were no regions where the regression accuracy was significantly higher in this time range.

**Fig. 9.**
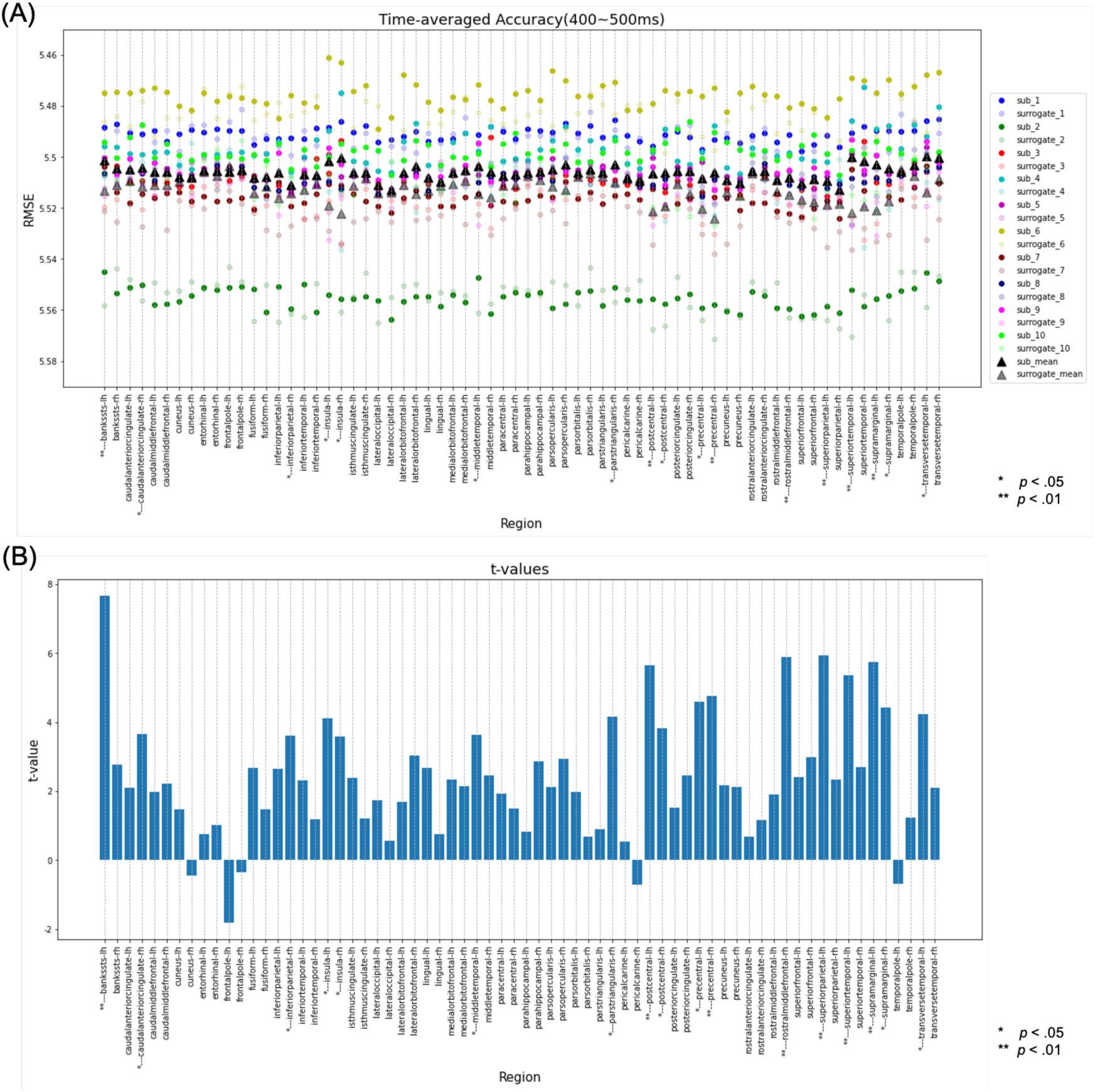
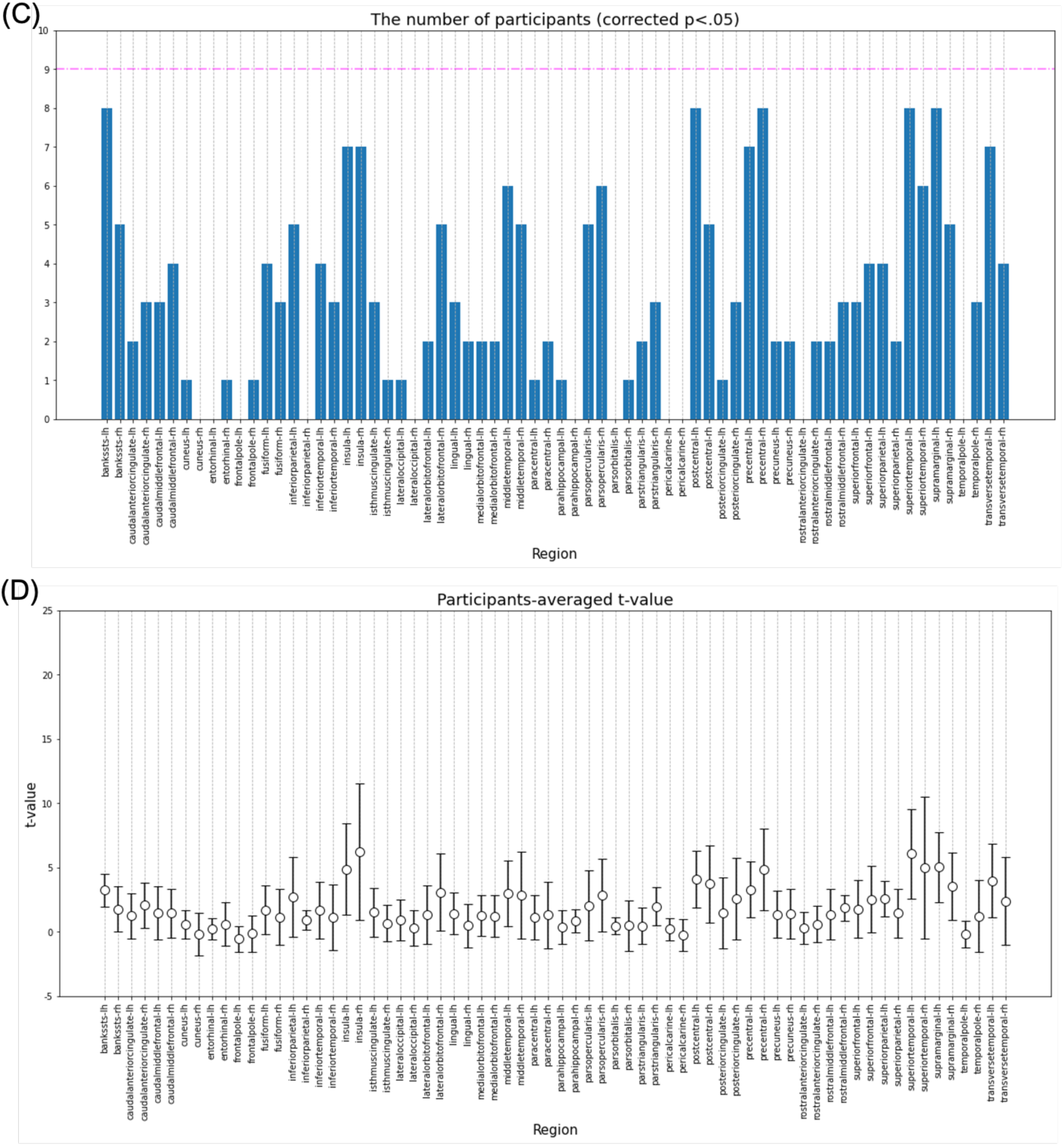
Results of the decoding analysis for each region in the time range of 400‒ 500 ms. (A) The regression error of the original data and the mean regression error of the surrogate data in the time range of 400‒500 ms were plotted for each subject. The paired-samples *t*-test for all subjects compared whether the regression error of the original data was significantly smaller than the mean regression error of the surrogate data. *Regions with a corrected *p*-value of < .05 and ** regions with a corrected *p*-value of < .01. (B) The *t*-value for each region in the paired-samples *t*-test. (C) Number of subjects with a corrected *p*-value of < .05 in the one-sample *t*-test comparing whether the regression error of the original data was significantly smaller than the regression error of the surrogate data for each subject. Regions where more than 90% of the total subjects (above the pink line) had a corrected *p*-value of < .05 were defined as regions with significant differences in the one-sample *t*-test. (D) Subject mean and variance of *t*-values in the one-sample *t*-test.

**Fig. 10.**
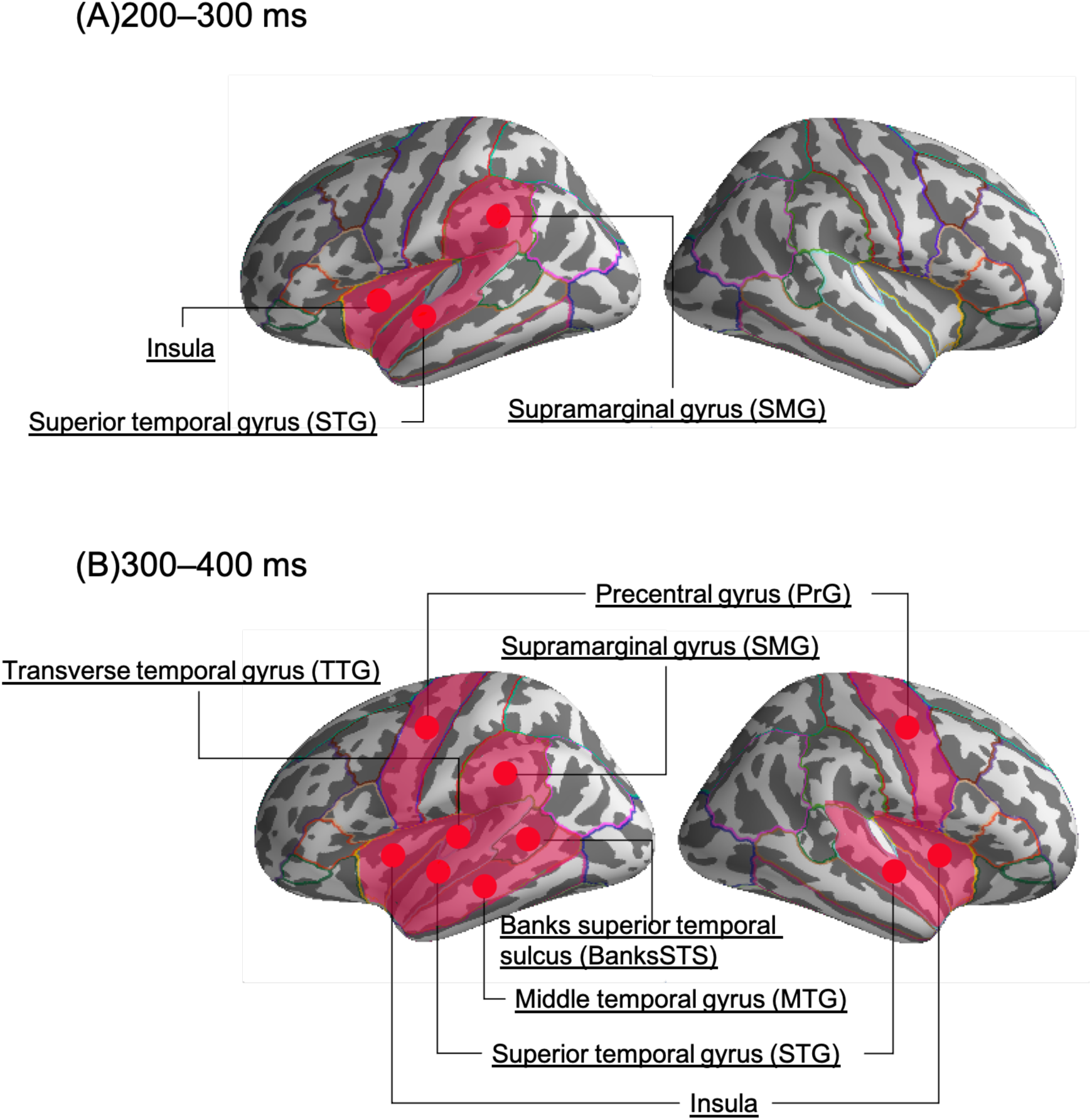
Regions where significant differences were found using both the paired-samples *t*-test and the one-sample *t*-test; that is, regions where the regression accuracy by decoding was determined to be significantly higher than the regression accuracy by surrogate. (A) Regions with significance at 200‒300 ms after the start of a long unit word (LUW). (B) Regions with significance at 300‒ 400 ms after the start of a LUW.

Figure 10 shows the regions with significantly higher regression accuracies based on the decoding analysis for each time range.

## 4. Discussion

In order to identify the brain mechanisms involved in human language prediction, this study measured brain activity during Japanese-language speech listening using MEG and examined its relationship with the surprisal predicted using the LSTMLM. For this study, we used two types of analyses: encoding and decoding. Some regions demonstrated significant differences using both types of analyses, including the left insula, left STG, and left SMG. On the other hand, some regions showed significant differences using either only encoding or decoding analyses (Table 1). In the encoding analysis, the left temporal pole, left FuG, bilateral IFG, and bilateral FOP showed significant differences. For the decoding analysis, the right insula, right STG, bilateral PrG, left TTG, left MTG, and left BanksSTS showed significant differences.

**Table 1.** Regions where significant differences were found in the encoding and/or decoding analyses. For the encoding analysis, regions with a corrected *p*-value of < .001 are represented as **, and regions with a corrected *p*-value of < .005 are represented as *. For the decoding analysis, regions where more than 90% of subjects had a corrected *p*-value of < .05 on the one-sample *t*-test and where there were significant differences in the paired-samples *t*-test, a corrected *p*-value of < .01 in the paired-samples *t*-test is indicated as **, and regions with a corrected *p*-value of < .05 are indicated as *. All regions determined to have significant differences using both types of statistical analyses in the 200‒300-ms time range were also included in the 300‒400-ms time range. For the 400‒500-ms time range, there were no regions where both statistics showed significant differences. Therefore, this table only shows the decoding results for the 300‒400-ms time range. BanksSTS, Banks superior temporal gyrus; FOP, frontal operculum; FuG, fusiform gyrus; IFG, inferior frontal gyrus; MTG, middle temporal gyrus; PrG, precentral gyrus; SMG, supramarginal gyrus; STG, superior temporal gyrus; TTG, transverse temporal gyrus

| Region |  | Encoding | Decoding (300–400 ms) |
| --- | --- | --- | --- |
| Insula | left | ** | ** |
|  | right | - | ** |
| STG | left | ** | ** |
|  | right | - | * |
| SMG | left | ** | ** |
|  | right | - | - |
| PrG | left | - | ** |
|  | right | - | ** |
| TTG | left | - | ** |
|  | right | - | - |
| MTG | left | - | ** |
|  | right | - | - |
| BanksSTS | left | - | ** |
|  | right | - | - |
| FuG | left | ** | - |
|  | right | - | - |
| IFG | left | * | - |
|  | right | * | - |
| FOP | left | * | - |
|  | right | * | - |
| Temporal pole | left | ** | - |
|  | right | - | - |

The regions that showed significant differences using both types of analyses were the left insula, the left STG, and the left SMG. These regions showed positive correlations between surprisal values and MEG source signals in the encoding analysis and also showed significantly higher regression accuracies at 200‒300 ms and 300‒400 ms after the onset of a LUW in the decoding analysis.

The left SMG is believed to be involved in comprehension of all types of language stimuli, including reading and listening. Previous studies have shown that the left SMG is activated more significantly when listening to or reading a story than when experiencing a nonverbal control stimulus (Bemis and Pylkkänen, 2013; Lindenberg and Scheef, 2007). The left SMG is also associated with phonological processing and has been shown to facilitate auditory memory and enable longer-term memory of musical pitches through stimulation and activation of the transcranial direct-current stimulation (tDCS) (Schaal et al., 2017). Therefore, the increased activity identified in the left SMG in this study may be due to the increased information-processing costs associated with language comprehension caused by the appearance of unpredicted LUWs.

The STG is composed of the primary and secondary auditory cortices and is believed to be responsible for phonological processing associated with speech understanding (Hickok and Poeppel, 2000). In particular, the left STG, which is a language-related area, is believed to be responsible for auditory short-term memory, language comprehension, and phonological processing (Buchsbaum et al., 2001; Leff et al., 2009). In addition, there have been positive correlations identified between STG activity measured by fMRI and the surprisal calculated by the N-gram language model during story listening. In a previous study, the STG’s activity was modulated by how well input words fit their predictions (Willems et al., 2016), and our results replicated the findings of this previous study.

The insula has been shown to be involved in several important processes, including auditory attention allocation, auditory adjustments to new stimuli, temporal processing, phonological processing, and auditory-visual integration. The left insula, in particular, is more involved in phonological word processing (Bamiou et al., 2003). Bilateral insula have been reported to be activated when listening to sentences that are not semantically valid (i.e., some of the words in the sentence are changed to inappropriate words), thus supporting semantic information processing at the sentence level (Friederici et al., 2003). Bilateral insula are also associated with risk and reward prediction errors in gambling tasks (Preuschoff et al., 2008). Therefore, the activation of the left insula identified in this study may have been caused by increases in phonological and semantic processing loads caused by the appearance of unexpected LUWs or by the encoding of prediction errors.

Some regions only showed significant differences in the encoding analysis (i.e., a significant correlation between surprisal and MEG source signals), including the left temporal pole, left FuG, bilateral IFG, and bilateral FOP. In particular, the left temporal pole and the left FuG showed positive correlations between surprisal and MEG source signals. The temporal pole region performs social and emotional processing, and the left temporal pole, in particular, has been closely associated with semantic memory (Olson et al., 2007). The left FuG is called the visual word form area (VWFA) (Cohen et al., 2000) and is responsible for the perception of visual word forms (Vinckier et al., 2007).

In addition, the inferior temporal sulcus, which is adjacent to the VWFA, has been shown to be involved in both visual and auditory word form comprehension (Cohen et al., 2004). The left temporal pole, left FuG, and STG have been shown to be positively correlated with brain activity during story listening measured on fMRI and the surprisal calculated by the N-gram language model. Our results replicate the findings of this previous study.

In the present study, bilateral IFG and FOP showed negative correlations between surprisal and MEG source signals. The IFG is responsible for syntactic processing and performs computations to predict grammatical structures (Friederici et al., 2006). The IFG has also been shown to be involved in sentence-level semantic processing and language-related working memory, with this region being significantly activated when listening to sentences during a task in which participants had to judge whether successive sentences had the same meaning (Dapretto and Bookheimer, 1999; Friederici, 2002). The FOP evaluates input elements for the grammatical structure predicted by the IFG (Friederici et al., 2006). Furthermore, this region is related to semantic processing of words, and its activity becomes stronger based on the weakness of the connections between adjacent words (Friederici, 2020). The relationship between surprisal and activity in the regions identified in this study is probably related to the fact that the surprisal can account for both semantic information at the sentence level and information on the connections between neighboring words when using the LSTMLM.

Some regions only showed significant differences in the decoding analysis (i.e., the original prediction accuracy was significantly higher than the surrogate prediction accuracy), including the right insula, right STG, bilateral PrG, left TTG, left MTG, and left BanksSTS. In a previous MEG study, the left BanksSTS and left MTG were shown to be significantly more activated during a visual reading task when reading a story than when reading a random word list. Thus, these regions are more strongly related to long-term cognitive processing at the sentence level than to short-term cognitive processing at the word level (Brennan and Pylkkänen, 2012). Although bilateral PrG are classified as part of the primary motor cortex, their activity has been shown to vary in relation to speech perception (Pulvermüller et al., 2006). The left TTG is classified as the primary auditory cortex and is implicated in pitch processing and in the learning of verbal pitch (Wong et al., 2008). The insula and STG, regions where significant left hemispheric differences were identified using both types of analyses in this study, also showed significant differences in the right hemisphere in the decoding analysis. The right insula and the right STG play similar roles to their counterparts in the left hemisphere by supporting semantic information processing at the sentence level (Friederici et al., 2003) and by phonological processing associated with speech understanding (Hickok and Poeppel, 2000).

Thus far, encoding analysis has more commonly been used in linguistic research than decoding analysis. Therefore, identification of the relationships between surprisal and brain activity in the additional regions identified in this study is an advantage of using decoding analysis. However, because the LSTMLM was used to estimate surprisal in this study, we cannot simply compare our results with previous studies that used the N-gram language model to estimate surprisal. Therefore, we cannot definitively determine whether the observed differences were an effect of the LSTMLM or the decoding analysis. Since the decoding analysis was performed independently for each region in this study, it is clear that information related to surprisal was processed as activity patterns in each of these regions of the brain. However, we did not examine the functional connectivity between regions. Human cognitive function is not the result of independent activity in a single brain region but rather the functional connectivity of multiple regions as a network. (Bressler and Menon, 2010). In order to clarify whether these new regions are directly related to surprisal or are secondary activities, it will be necessary to examine how these regions are connected to other surprisal-related regions and how they function as networks.

In terms of the temporal response in brain activity, the encoding analysis showed an increase in activity, with a peak at 300 ms after the onset of a LUW. The decoding analysis showed higher prediction accuracies of a larger number of regions in the 300‒400-ms time range than in the 200‒300-ms time range, with no regions showing significant decoding accuracies at 400‒500 ms after the onset of a LUW. It is possible that this temporal response pattern corresponds with the N400, a negative brain activity measured on the scalp at about 400 ms after a linguistic stimulus. The N400 is known to have access to semantic memory (Kutas and Federmeier, 2000). It has also been shown that the magnitude of the surprisal correlates with the amplitude of the N400 measured on the scalp using EEG (Fitz and Chang, 2019). In addition, the N400 observed in MEG source signals has been shown to originate from the left STG at approximately 250 ms after the onset of stimulation and to gradually spread (Kutas and Federmeier, 2011). In this study’s decoding analysis, significant differences were found in a small number of regions in the left hemisphere in the time range of 200‒300 ms, while significant differences were found in a broader range of regions, including the right hemisphere, in the 300‒400-ms time range. This finding implies that the N400 occurred in the left hemisphere and spread to a broader region.

We believe the language model used in this study was relatively valid because significant differences were found in regions suggested by previous studies. Moreover, the N-gram language model was not able to learn surprisal values using data from this study, while the LSTMLM was able to learn the surprisal values to some extent. Therefore, the LSTMLM was able to estimate the surprisal with a smaller amount of data, demonstrating its effectiveness at performing these estimations. However, the dataset for this study was small, even for the LSTMLM, and it is therefore difficult to claim that this model was able to achieve sufficient accuracy. Huge data sets, such as sentences from news articles, are generally used in language-model training; however, we did not use these types of datasets in this study because the speeches used in this experiment were spoken, and, therefore, training with a written language dataset could have resulted in a wording bias. Also, since humans generally listen to spoken language in their daily lives, there should be more spoken language than written language in training data. Therefore, a large dataset of spoken Japanese should be used as training data for models designed to represent language processing in the human brain; however, the dataset used in the present study is currently the largest one available. In order to clarify the relationship between surprisal and brain activity, future studies should enrich the spoken language dataset for training data and should develop language models that can be trained with smaller amounts of data. However, in studies of the human brain, it is important to avoid language models that ignore the structure of the human brain even if they do improve the accuracy of the language model because these types of models cannot accurately represent language processing.

This study had several limitations. First, the number of participants was only 10, which is a smaller sample size than in similar MEG studies (Armeni et al., 2019: 25 participants). Also, we used the FreeSurfer average brain to adjust the MEG sensor positions instead of MRI structural images for each participant. Therefore, we could not account for individual differences in brain structure, and the accuracy of the signal source estimation was therefore questionable. Next, in the decoding analysis, we performed Lasso regression for each region; however, we did not examine which sources were selected as features after regularization and how many of them were used in each region. In addition, the number of sources contained in each pre-regularized region were different. Therefore, we cannot exclude the possibility that the regression accuracy was affected by too many or too few features in each region.

In future studies, it will be necessary to compare language models with identical experimental paradigms to clarify whether the results of this study were the effect of the LSTMLM or the decoding analysis. In addition, it will be necessary to understand the broader network of the language prediction mechanism in order to investigate the functional connectivity between regions and not just in individual regions.

## 5. Conclusion

In order to investigate the brain mechanisms involved in human language prediction, this study clarified the relationship between the surprisal estimated by the LSTMLM and MEG source signals while listening to Japanese speech using encoding and decoding analyses. In addition to surprisal-related regions revealed in previous studies, such as the STG, FuG, and temporal pole, we also identified relationships between surprisal and brain activity in other regions, including the insula, STS, and MTG, which are believed to be engaged in longer-term, sentence-level cognitive processing.

## Notes

### Competing Interest Statement

The authors have declared no competing interest.

